# Kynurenine pathway metabolomics in heatstroke: a validated LC-MS/MS method reveals compartment-specific neurochemical disruption in a murine model

**DOI:** 10.64898/2026.06.29.735282

**Authors:** Petra Majerova, Dorothy Wasike, Juraj Piestansky, Andrej Kovac

## Abstract

Heat stroke is characterized by profound central nervous system dysfunction and vascular abnormalities. Previous studies have demonstrated the marked vulnerability of the CNS to thermal stress, resulting in neuronal injury and glial activation. However, the metabolic mechanisms linking acute injury to chronic neurological long-term effects remain understood. The neuropathological changes are closely associated with neuroinflammatory and metabolic disturbances, including dysregulation of the kynurenine pathway, whose metabolites modulate neurotoxicity, neuroprotection, and immune responses. Here, we present the first comprehensive characterization of kynurenine pathway metabolomic profile across both plasma and brain tissue in a mouse model of heat stroke. Using a validated and sensitive LC-MS/MS method, we simultaneously measured and quantified 13 analytes (kynurenine, kynurenic acid, quinolinic acid, nicotinic acid, picolinic acid, xanthurenic acid, anthranilic acid, 3-hydroxykynurenine, 3-hydroxyanthranilic acid, indole-3-acetic acid, indole-3-lactic acid, 5-hydroxyindoleacetic acid and neopterin). The findings reveal a biphasic metabolic response, characterized by an acute serotonergic disruption and reduced neuroprotective capacity, followed by chronic activation of the kynurenine pathway, depletion of central serotonin metabolites, and metabolic signatures consistent with gut microbiota dysbiosis. The acute phase is marked by a transient imbalance favoring neurotoxic kynurenine pathway metabolites, whereas the chronic phase reflects sustained pathway activation. Notably, the plasma-brain dissociation of 5-hydroxyindoleacetic acid emerged as the most prominent cross-compartment finding, suggesting a potential biomarker of central serotonergic depletion and a mechanistic link between peripheral and central metabolic changes, with implications for therapeutic targeting during the subacute recovery phase.

## INTRODUCTION

Climate change has contributed to rising global temperatures and an increased frequency and intensity of heatwaves. Consequently, the incidence of heat-related illnesses (HRIs), ranging from mild heat exhaustion to life-threatening heatstroke, has also increased, as demonstrated by nationwide epidemiological studies reporting a progressive rise in HRI incidence over recent decades^1–3^. Heatstroke (HS) is the most severe and life-threatening form of heat injury. The growing burden of extreme heat exposure underscores the need to better understand the pathophysiological consequences of HS, particularly in vulnerable populations-elderly and children.

Clinically, HS is characterized by CNS dysfunction, vascular abnormalities, and a pronounced neuroinflammatory activation. Previous studies have demonstrated heat-induced neuronal damage and activation of astrocytes and other neuroglial cells, as evidenced by increased expression of glial markers and neuroinflammatory changes in vulnerable brain regions, including the hippocampus and cerebellum^4,5^. Experimental findings from a mouse model designed to replicate clinically relevant heat exposure conditions revealed cerebellar demyelination and Purkinje cell loss^6^. In addition, hyperthermia has been shown to promote oxidative stress, mitochondrial dysfunction, and disruption of the blood-brain barrier (BBB), thereby contributing to neurological dysfunction following HS^7,8^.

Accumulating evidence indicates that oxidative stress, neuronal damage, and inflammatory responses are accompanied by significant alterations in kynurenine pathway (KP) metabolism. KP is the primary route for tryptophan (TRP) degradation. TRP is an essential amino acid, and approximately 90-95% of its degradation occurs through the KP, with the liver accounting for nearly 90% of total TRP catabolism^9^. Although under normal physiological conditions only a small proportion of TRP is converted into its central metabolite, kynurenine (KYN) (∼5-10%), this conversion is markedly increased during inflammation^10–12^. TRP degradation to KYN is catalyzed by indoleamine 2,3-dioxygenase (IDO), which is strongly upregulated by pro-inflammatory cytokines, particularly interferon-γ (IFN-γ)^13,14^. IFN-γ also induces neopterin production in activated monocytes, making neopterin a well-established biomarker of cellular immune activation^15^.

KYN is further metabolized into a range of downstream KP metabolites through the action of key enzymes, including kynurenine monooxygenase, kynureninase, and kynurenine aminotransferases. These metabolites possess diverse immunomodulatory and neuroactive properties. Kynurenic acid (KYNA) is generally considered neuroprotective. KYNA exerts anti-glutamatergic effects by acting as an antagonist of the N-methyl-D-aspartate (NMDA) receptor, thereby reducing glutamate-mediated excitotoxicity and conferring neuroprotective effects in experimental models^16–18^. Quinolinic acid (QA), picolinic acid (PA) and nicotinic acid (NA) are end products of the KP and serve as intermediates in the biosynthesis of nicotinamide adenine dinucleotide (NAD⁺)^19^. These compounds are involved in complex interactions with inflammatory and apoptotic processes that contribute to neuronal injury and cell death in the CNS^20,21^.

In addition, KP metabolites exhibit distinct redox properties. KYNA and 3-hydroxyanthranilic acid (3OH AA) exert antioxidant effects through free radical scavenging and inhibition of lipid peroxidation, whereas 3-hydroxykynurenine (3OH KYN) promotes oxidative stress via reactive oxygen species generation^22,23^. Furthermore, both KYN and 3OH AA have been shown to exert immunosuppressive properties, including the induction of regulatory T-cell responses and the suppression of effector immune cell proliferation and survival^24,25^. This suggests that the KP is an important modulator of inflammatory and neuroimmune processes. Through combined redox and immunomodulatory properties, KP metabolites may contribute to the regulation of oxidative stress associated with neuronal damage in CNS.

There are limited studies investigating the KP in heatstroke. Previous experimental work has shown that KYNA administration significantly prolongs survival time and attenuates systemic and neural injury in rat models of HS. Vehicle-treated HS rats exhibited hypotension, hypothalamic neuronal degeneration and apoptosis, elevated serum levels of tumor necrosis factor-α (TNF-α), intercellular adhesion molecule-1 (ICAM-1), and interleukin-10 (IL-10), as well as multiorgan apoptosis^26^. In contrast, KYNA preconditioning attenuated hypotension and hypothalamic neuronal injury, reduced apoptosis in peripheral organs, increased IL-10, and decreased TNF-α and ICAM-1 levels, indicating modulation of inflammatory and apoptotic responses during HS.

A human study comparing exertional and exogenous heating reported increased plasma KP metabolites, including KYNA, PA, 3OH KYN, and QA, alongside elevated stress markers in both conditions; however, these changes were transient and returned to baseline during recovery^27^.

Here, we present the first comprehensive characterization of the KP metabolomic profile in both plasma and brain tissue in a mouse model of HS. We simultaneously quantified 13 kynurenine pathway metabolites across acute and long-term time points (1 week and 3 weeks post-HS), and assessed whether their levels return to baseline during recovery. Furthermore, we investigated whether HS-induced alterations in kynurenine metabolism preferentially shift toward neurotoxic or neuroprotective pathways, providing insight into their potential contribution to heatstroke-associated neurodegeneration.

## MATERIALS AND METHODS

### Experimental animals

Male C57BL/6J mice aged 20-24 weeks were used in this study. All animals were bred in the in-house animal facility of the Institute of Neuroimmunology of the Slovak Academy of Sciences in Bratislava. All animals were housed under standard laboratory conditions with ad libitum access to food and water. Mice were maintained on a 12-h light/dark cycle (lights on at 7:00 a.m.) at room temperature (24 ± 2 °C) and constant relative humidity (40 ± 15%). All experiments on animals were carried out according to the institutional animal care guide-lines conforming to international standards (Directive 2010/63/EU) and were approved by the State Veterinary and Food Committee of Slovak Republic (RO-1101/14-221C). Efforts were made to minimize the number of animals utilized and limit discomfort, pain, or any other suffering of the experimental animals in this study.

### Chemicals and Reagents

Analytical standards of all analytes were purchased from Sigma Aldrich (Steinheim, Germany). Isotopically labelled internal standards (IS) were obtained as follows: PA-d4, NA-d4 and 3-indole-3-acetic acid (IAA-d7) from Merck (Darmstadt, Germany); Kynurenine-d4, KYNA-d5 from CDN isotopes (Pointe Claire, Quebec, Canada), xanthurenic acid (XA-d4) from Santa Cruz Biotechnology (Dallas, Texas, USA), 3OH KYN-d3 from Buchem BV (Minden, Apeldoorn, Netherlands), QA-d3 and 5-hydroxyindoleacetic acid (5OH IAA-d6) from MedChemTronica (Sollentuna, Sweden); 3OH AA-d2 and indole-3-lactic acid (I3LA-d5) from LGC Standards (Łomianki, Poland), anthranilic acid (AA-ring-13C6) from Eurisotope (Saint-Aubin, France), Neopterin 13C5 from MedChemExpress (Scintila, s.r.o., Slovakia). Ammonium formate (AF), formic acid (FA) for LC-MS were obtained from Sigma Aldrich. LC-MS grade methanol (MeOH), acetonitrile (ACN) and water were acquired from Merck (Seelze, Germany).

### Standards and Reagents Preparation

Stock standard solutions of all analytes were prepared in DMSO, in concentration 1 mg/mL, aliquoted and stored under -80°C for further use. Isotopically labeled internal standards were prepared in the same manner, aliquoted and stored under -80 °C for further use. Standards and IS were then diluted with initial mobile phase as needed.

### Heat exposure protocol

Experimental parameters to ensure heatstroke induction were validated in previous study^28^. To investigate the pathogenesis of heatstroke, a murine heatstroke model was induced by elevating the animals’ core body temperature to 41 °C. A semi-enclosed heatstroke chamber (200 × 340 × 300 mm) made of acrylic was constructed by vertically stacking animal cages in a greenhouse-like configuration. An ultrasonic humidifier (SMB-1, Sakura Seiki, Tokyo, Japan) and a digital thermo-hygrometer (AD-5696, A&D Company, Tokyo, Japan) were used to control and monitor ambient temperature (AT), relative humidity (RH), and wet-bulb globe temperature (WBGT). Prior to the experiment, the heatstroke chamber, humidifier, thermo-hygrometers as well as a camera, an oxygen pump, and light sources were pre-warmed inside an incubator overnight at 30 °C. The heatstroke chamber was preheated to the desired experimental temperature for at least 3 h. The humidifier was started 3 h before heat exposure to establish a hot and humid environment. During this period, mice were subjected to 3 h of water restriction. Mildly dehydrated mice were then placed in the heatstroke chamber, exposed to heat (AT 41 °C, RH > 95%) for 60 min, and subsequently returned to their home cages maintained at room temperature (Fig. 1). Body weight (BW) was measured as an indicator of body fluid volume: prior to water restriction, after 3 h of water restriction and immediately after heat exposure. After heatstroke (1 and 3 weeks), cerebellum and plasma were collected. The brain tissue was frozen at −80 °C for subsequent analyses. Blood was collected into EDTA tubes and centrifuged for 10 min at 2,000 × g. Plasma aliquots (50 µL) were prepared and stored at −80 °C for further analysis.

**Figure 1.**
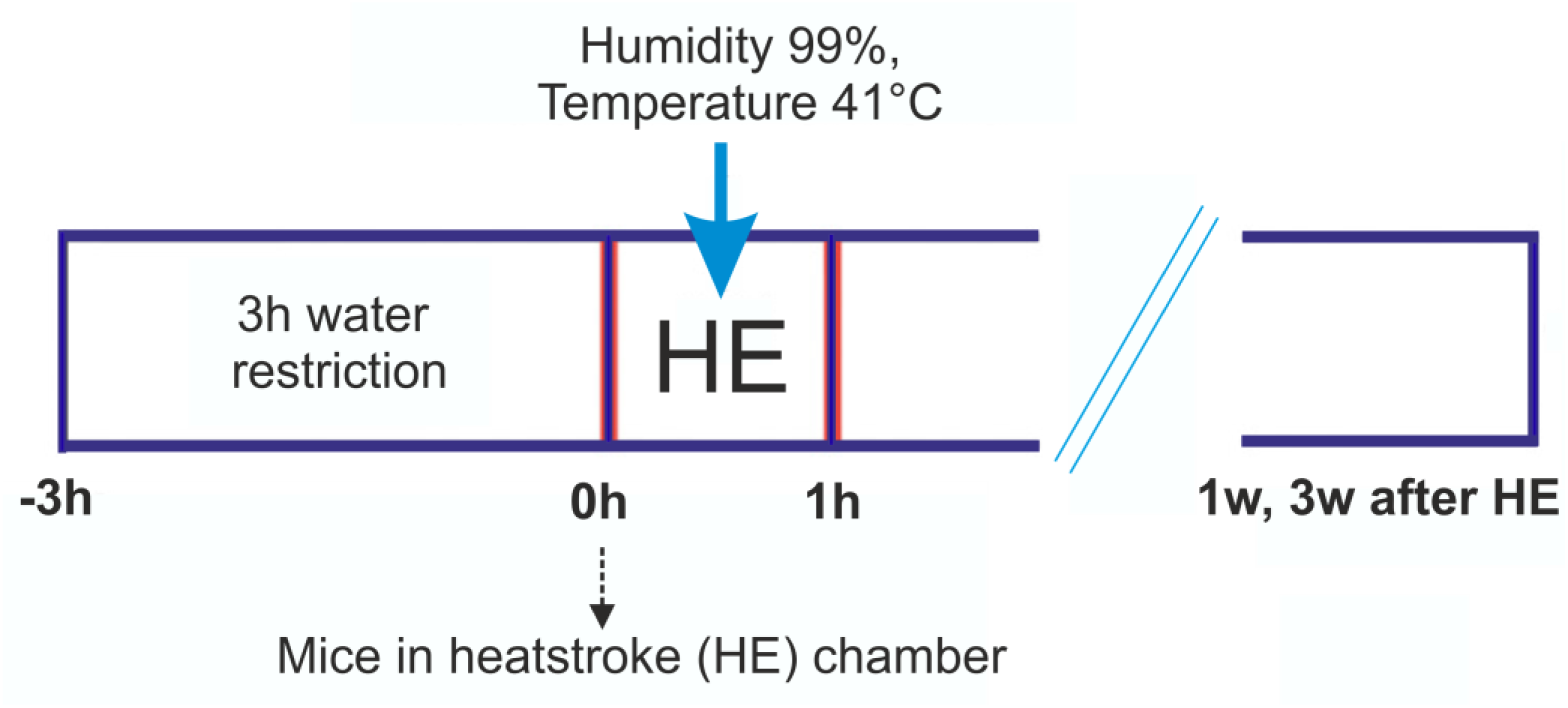
Experimental design. Heatstroke model was induced by elevating the animals’ core body temperature to 41 °C. At the selected time points plasma and brain tissue was collected and snap frozen in liquid nitrogen.

### Plasma and brain tissue collection

Plasma was obtained from control mice and from mice at 24, 48 hours, 1 and 3 weeks after HS exposure. Plasma was separated by centrifugation at 3,000 rpm for 10 minutes and stored at −80 °C until analysis. Brain tissue was briefly perfused with cold PBS/heparine and dissected. Tissue samples were snap-frozen in liquid nitrogen and stored at −80 °C until analysis.

### Preparation of samples for LC/MS analysis

Plasma samples for quantification were prepared as follows. First, 2.5 µL of internal standards were added to LC-MS-grade methanol (Honeywell, North Carolina, USA). Then, 50 µL of plasma was precipitated by the addition of 200 µL of methanol containing the standards. The use of stable isotope-labelled internal standards for each analyte class ensures accurate quantification despite matrix-specific ion suppression. Samples were vortexed and centrifuged at 30,000 × g for 10 min at 4 °C. The supernatants were transferred to a deep-well plate and dried under vacuum. The residues were reconstituted in initial mobile phase, centrifuged again at 30,000 × g for 10 min at 4 °C, and transferred into vials (Waters, Milford, MA, USA) for LC-MS analysis. Approximately 20 mg of brain tissue was weighed and homogenized in water (150 µL per 20 mg tissue) using stainless steel beads (SSB16, Next Advance, USA) in a FastPrep-24™ instrument (MP Biomedicals, USA) at a speed of 4.5 m s⁻¹ for 20 s. The homogenates were then centrifuged at 30,000 × g for 10 min at 4 °C. Subsequently, 100 µL of supernatant was transferred to a new tube, and 400 µL methanol containing internal standards was added for protein precipitation. After centrifugation (30,000 × g for 10 min at 4 °C), 450 µL of the supernatant was evaporated using a Savant SpeedVac SPD111V (Thermo Fisher Scientific, USA). The samples were reconstituted in initial mobile phase, centrifuged again, transferred into vials, and subjected to LC-MS analysis.

### Ultra-performance liquid chromatography coupled to tandem mass spectrometry

All analyses were performed on ACQUITY UPLC® H class Premier system (Waters, Milford, MA, USA) consisting of solvent manager, sample manager flow-through needle (FTN), and column manager equipped with reversed-phase column Waters ACQUITY UPLC® HSS T3 1.8 μm (2.1×150 mm) and ACQUITY UPLC® HSS T3 1.8 μm (2.1×5 mm) VanGuard pre-column. The mobile phase consisting of 0.2% FA in 30mM ammonium formate buffer (A) and 0.2% FA in ACN (B). For analytes separation, gradient elution, with total analysis time of 8 min, was used: starting at 1% of B, increasing to 10% (0-1.5min), then to 60 % (1.5-3.5 min), increasing to 90% (3.5-4.5) hold for 0.5 min (4.5-5 min) and return to initial conditions in 0.5 min (5-5.5 min) and re-equilibrating for 2.5 min (5.5-8.0 min). The flow rate of the mobile phase was 0.4 mL/min, the column temperature was 30 °C, and the sample injection volume was 5 µL. The LC system was connected with triple quadrupole mass spectrometer Xevo TQ-Absolute (Waters, Milford, MA, USA) equipped with electrospray ionization source (ESI) working in positive mode. ESI+ conditions were as follows: cone gas flow 150 L/h, desolvation gas flow 900 L/h, source temperature 150 °C, desolvation temperature 550 °C, capillary voltage 3 kV. Collision energies and source cone voltage were manually tuned for each analyte and corresponding IS. ESI-conditions were as follows: cone gas flow 50 L/h, desolvation gas flow 900 L/h, source temperature 150 °C, desolvation temperature 550 °C, capillary voltage 2 kV. Collision energies and source cone voltage were manually tuned for each analyte and corresponding IS. LC-MS system was controlled, and all data were acquired by MassLynx V4.2 software (Waters, Milford, MA, USA).

### Statistical analysis

Differences among the experimental groups were evaluated using the Kruskal-Wallis one-way analysis of variance by ranks. Where the Kruskal-Wallis test indicated a significant overall effect (p < 0.05), pairwise post-hoc comparisons were performed using Dunn’s test with Bonferroni correction for multiple comparisons. Corrected p-values are reported for all pairwise comparisons; a corrected p-value of < 0.05 was considered statistically significant. All statistical analyses were performed in Python (version 3.12) using the SciPy library (version 1.13).

## RESULTS

### Development and optimization of UPLC/MS method

NA and PA are isomeric compounds with identical MS/MS transitions, so chromatographic resolution is essential to distinguish them. Moreover, many KP metabolites occur at low basal concentrations in plasma and brain. UHPLC conditions were therefore optimized to achieve optimal chromatographic separation and to achieve maximum sensitivity of the mass spectrometry detection. The ionisation and fragmentation behaviours of all analytes were also studied. We used both positive and negative electrospray ionization since it easily ionize molecules with ionizable functional groups that are typical for KP metabolites. All analytes were detected by MS/MS under collision activated dissociation (CAD) conditions. Using the selected reaction monitoring (SRM) mode, fragment spectra were generated. For each analyte two transitions were measured, one was used for quantification, second as confirmatory. For the internal standards one transition was measured. The mass spectrometry parameters are summarized in Table 1.

**Table 1.** SRM conditions of metabolites and their internal standards.

| Analyte | Precursor ion | Product ion (qualitative) | Product ion (quantitative) | Cone [V] | Ce [eV] | ESI polarity |
| --- | --- | --- | --- | --- | --- | --- |
| QA | 166 | 122 | 77.95 | 20 | 15 | ESI- |
| QA-d3 | 169 | - | 81 | 20 | 15 | ESI- |
| NA | 124 | 53.03 | 80 | 30 | 20 | ESI+ |
| NA-d4 | 128 | - | 84.1 | 30 | 20 | ESI+ |
| PA | 124 | 106 | 77.8 | 20 | 15 | ESI+ |
| PA-d4 | 128 | - | 110 | 20 | 15 | ESI+ |
| Neopterin | 254 | 190 | 206 | 20 | 20 | ESI+ |
| Neopterin-13C5 | 259 | - | 210 | 20 | 20 | ESI+ |
| 3OH KYN | 225.1 | 161.9 | 208 | 20 | 20/10 | ESI+ |
| 3OH KYN-d3 | 228.1 | - | 111.1 | 20 | 20 | ESI+ |
| Kynurenine | 209.1 | 146.1 | 94.1 | 20 | 15 | ESI+ |
| Kynurenine-d4 | 213.1 | - | 98.1 | 20 | 15 | ESI+ |
| XA | 206 | 132 | 160 | 20 | 25/17 | ESI+ |
| XA-d4 | 210.1 | - | 164 | 20 | 17 | ESI+ |
| KYNA | 190 | 116 | 88.9 | 30 | 25 | ESI+ |
| KYNA-d5 | 195.1 | - | 121 | 30 | 25 | ESI+ |
| 3OH AA | 152 | 91 | 108 | 20 | 20/15 | ESI- |
| 3OH AA-d2 | 154.2 | - | 110 | 20 | 15 | ESI- |
| 5OH IAA | 192 | 91 | 146 | 30 | 30/15 | ESI+ |
| 5OH IAA-d6 | 198 | - | 151 | 30 | 15 | ESI+ |
| I3LA | 204 | 158 | 142 | 20 | 15/20 | ESI- |
| I3LA-d5 | 209 | - | 163 | 20 | 15 | ESI- |
| AA | 136 | 65.1 | 92 | 20 | 30/15 | ESI- |
| AA-d4 | 140 | - | 96.1 | 20 | 15 | ESI- |
| IAA | 176 | 103.1 | 130.1 | 20 | 15 | ESI+ |
| IAA-d7 | 183 | - | 137 | 20 | 15 | ESI+ |

KYNA, kynurenic acid; XA, xanthurenic acid; NA, nicotinic acid; PA, picolinic acid; AA, anthranilic acid; 3OH KYN, 3-hydroxykynurenine; 3OH AA, 3-hydroxyanthranilic acid; 5OH IAA, 5-hydroxyindole-3-acetic acid; QA, quinolinic acid; I3LA, indole-3-lactic acid; IAA, indole-3-acetic acid

Optimisation of the gradient programme and mobile phase composition demonstrated that 30 mM ammonium formate with 0.2% formic acid as the aqueous phase, paired with acetonitrile containing 0.2% formic acid as the organic modifier, provided optimal chromatographic resolution across all 13 analytes within an acceptable run time. The analytes and associated internal standards were detected at 1.26 min for QA, 1.49 min for PA, 1.64 min for neopterin, 1.65 min for NA, 2.01 min for 3OH KYN, 2.69 min for kynurenine, 2.91 min for KYNA, 3.1 min for 3OH AA, 3.24 min for 5HIAA, 3.63 min for I-3-LA, 3.72 min for AA and 3.92 min for IAA (Fig. 2).

**Figure 2.**
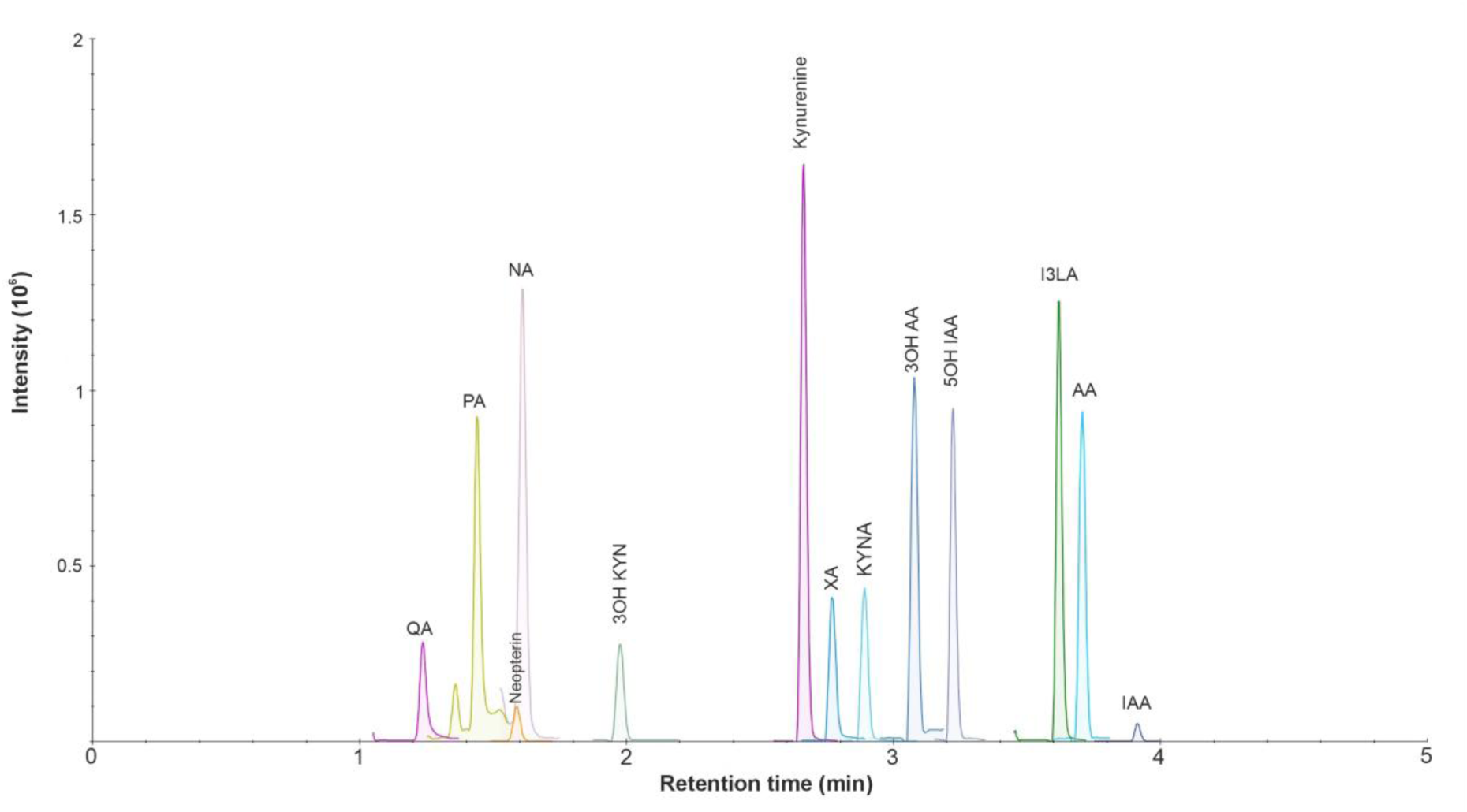
Extracted ion chromatograms of 13 targeted. Selected reaction monitoring (SRM) was used to detect targeted analytes. Chromatographic conditions: column, Acquity HSS T3 column (2.1 mm × 150 mm, 1.7 µm particle size) with VanGuard pre-column; flow rate = 0.4 mL min^-1^; column temperature 40 °C.

### Method validation

The method was validated for both matrixes in accordance with current acceptance criteria for bioanalytical method validation by European medicine Agency (EMA)^29^. Linear calibration curves (correlation coefficient > 0.99) were obtained for all 13 analytes within the concentration ranges studied for both matrixes. The linear regression equations, correlation coefficients, limits of detection and quantification are summarised in Supplementary Table S1-2. Precision was expressed as the relative coefficient of variation (CV%) of the within-run (intra-day) and between-run (inter-day) analytical results. The intra-day and inter-day precision and accuracy results are shown in Supplementary Table S3-7. In plasma, intra-day precision ranged from 0.6% to 14.5% CV and accuracy was within 87.1-113.1% across all four QC levels and 13 analytes. Inter-day precision (n = 18; three independent runs) ranged from 1.7% to 9.4% CV and accuracy was within 92.5-108.1%. In brain tissue, intra-day precision ranged from 1.5% to 12.4% CV and accuracy was within 88.8-113.2% across three QC levels (QC1-QC3). Inter-day precision ranged from 1.2% to 10.7% CV and accuracy was within 94.3-104.5%. All values were within the acceptance criteria of ±15% for accuracy and CV ≤15%, in accordance with EMA/FDA bioanalytical method validation guidelines.

Extraction recovery values ranging from 83% to 104% across the low and high QC levels. The matrix effects ranged from 92% to 116% (Supplementary Table S8). All of the analytes were stable in autosampler at 4°C for 24 h (Supplementary Table S9). All analytes were freeze-thaw stable over three cycles, except 3OH AA (Supplementary Table S10).

### Heatstroke induces delayed and sustained alternations of KP metabolites in plasma

To determine the impact of HS on systemic KP metabolism, plasma concentrations of KP metabolites were quantified immediately after HS and at 24 h, 48 h, 1 week, and 3 weeks post-insult. Comparison across post-HS time points revealed a distinct temporal pattern characterized by delayed and sustained alterations in selected KP metabolites rather than an immediate disruption of the pathway (Fig. 3).

**Figure 3.**
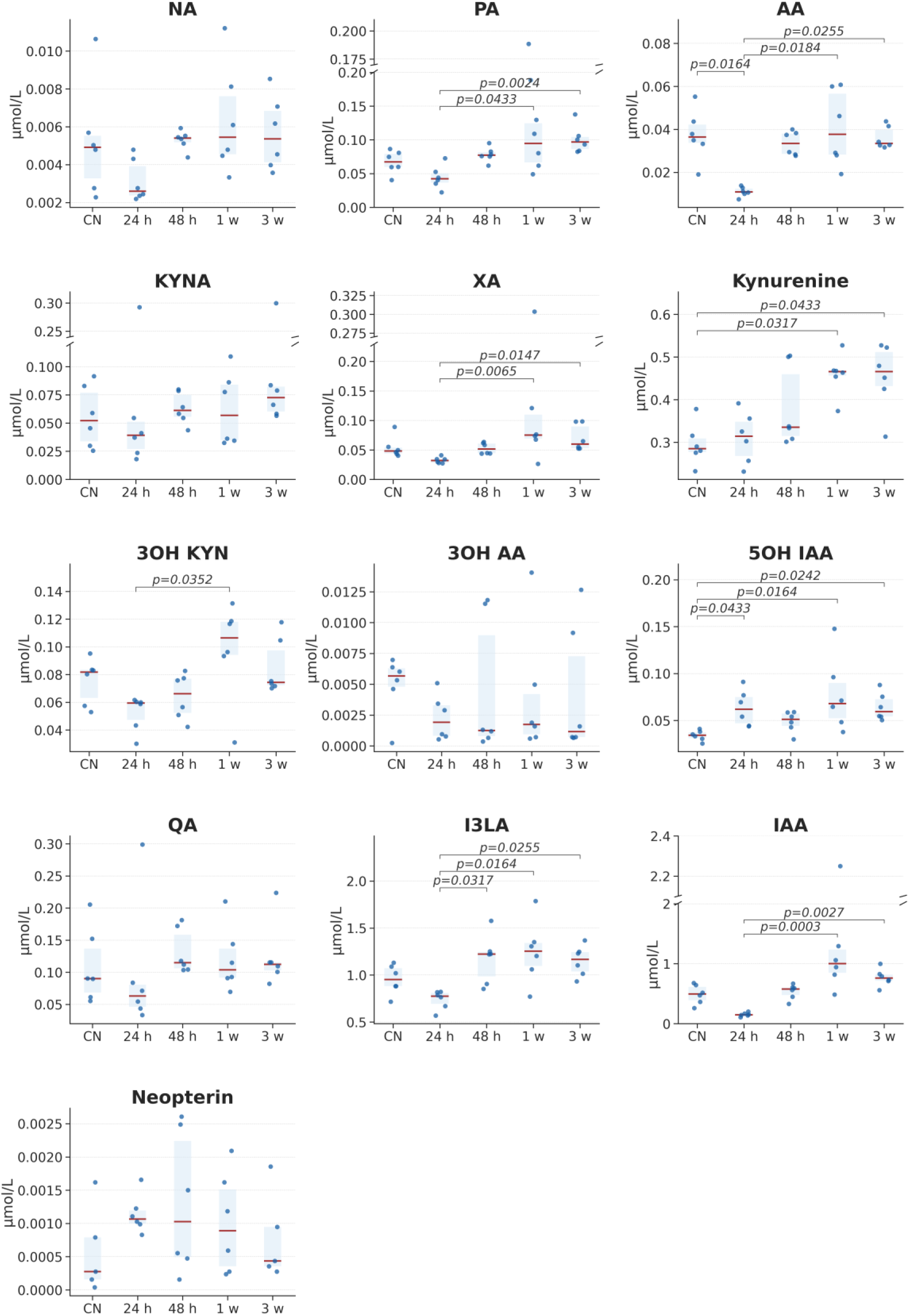
Changes of KP metabolites in plasma after the HS. Four different time points were analyzed, 24h, 48h, 1 week and 3 weeks. Boxes represent the interquartile range with median (red line); individual data points are shown. p-values indicate significant pairwise differences (Dunn’s test with Bonferroni correction following Kruskal-Wallis). CN, control (n = 6); 24 h, 48 h, 1 w, 3 w: time points post-heatstroke (n = 6 per group).

Among the metabolites analyzed, 5OH IAA was the only metabolite consistently elevated relative to control animals, showing significantly increased concentrations at 24 h (p = 0.0433), 1 week (p = 0.0164), and 3 weeks (p = 0.0242) after HS. In contrast, several KP metabolites displayed significant temporal changes within the HS group that emerged predominantly during the subacute and chronic phases of recovery.

Plasma 3OH-KYN increased at 1 week compared with 24 h post-HS (p = 0.0352), followed by a decline at 3 weeks, suggesting a transient activation of the oxidative branch of the KP. Similarly, I3LA exhibited progressive increases at 48 h (p = 0.0317), 1 week (p = 0.0164), and 3 weeks (p = 0.0255) relative to 24 h post-HS. A comparable time-dependent pattern was observed for IAA, with concentrations increasing markedly at 1 week (p = 0.0003) and 3 weeks (p = 0.0027) compared with 24 h post-HS.

XA also showed significant elevations at 1 week (p = 0.0065) and 3 weeks (p = 0.0147), indicating sustained activation of downstream kynurenine metabolism during recovery. AA was significantly reduced at 24 h post-HS compared with control animals (p = 0.0164), and remained lower than the 24 h value at 1 week (p = 0.0184) and 3 weeks (p = 0.0255).

PA, another downstream product of the pathway, was similarly increased at 1 week (p = 0.0433) and 3 weeks (p = 0.0024). These findings suggest prolonged activation of distal KP metabolism following HS. Although NA levels showed a tendency to increase after HS, no significant differences were detected compared with control animals.

Direct comparison with control animals revealed significantly elevated kynurenine concentrations at 1 week (p = 0.0317) and 3 weeks (p = 0.0433), whereas no significant differences were detected for the remaining time points. Furthermore, KYNA, 3OH AA, QA and neopterin remained unchanged throughout the experimental period.

Collectively, these data demonstrate that HS induces a selective and predominantly delayed remodeling of KP metabolism. The observed increases in kynurenine, 3OH KYN, XA, and PA during the subacute and chronic phases suggest sustained activation of downstream kynurenine metabolism that persists beyond the acute phase of thermal injury. The delayed nature of these alterations may reflect prolonged systemic inflammatory signaling, ongoing oxidative stress, and metabolic adaptation during recovery rather than an immediate response to the thermal insult itself.

### Heatstroke induces time-dependent alterations in KP metabolites in the brain

To evaluate the impact of HS on brain KP metabolism, metabolite concentrations were analyzed in the frontal cortex, a region particularly vulnerable to thermal injury (Fig. 4).

**Figure 4.**
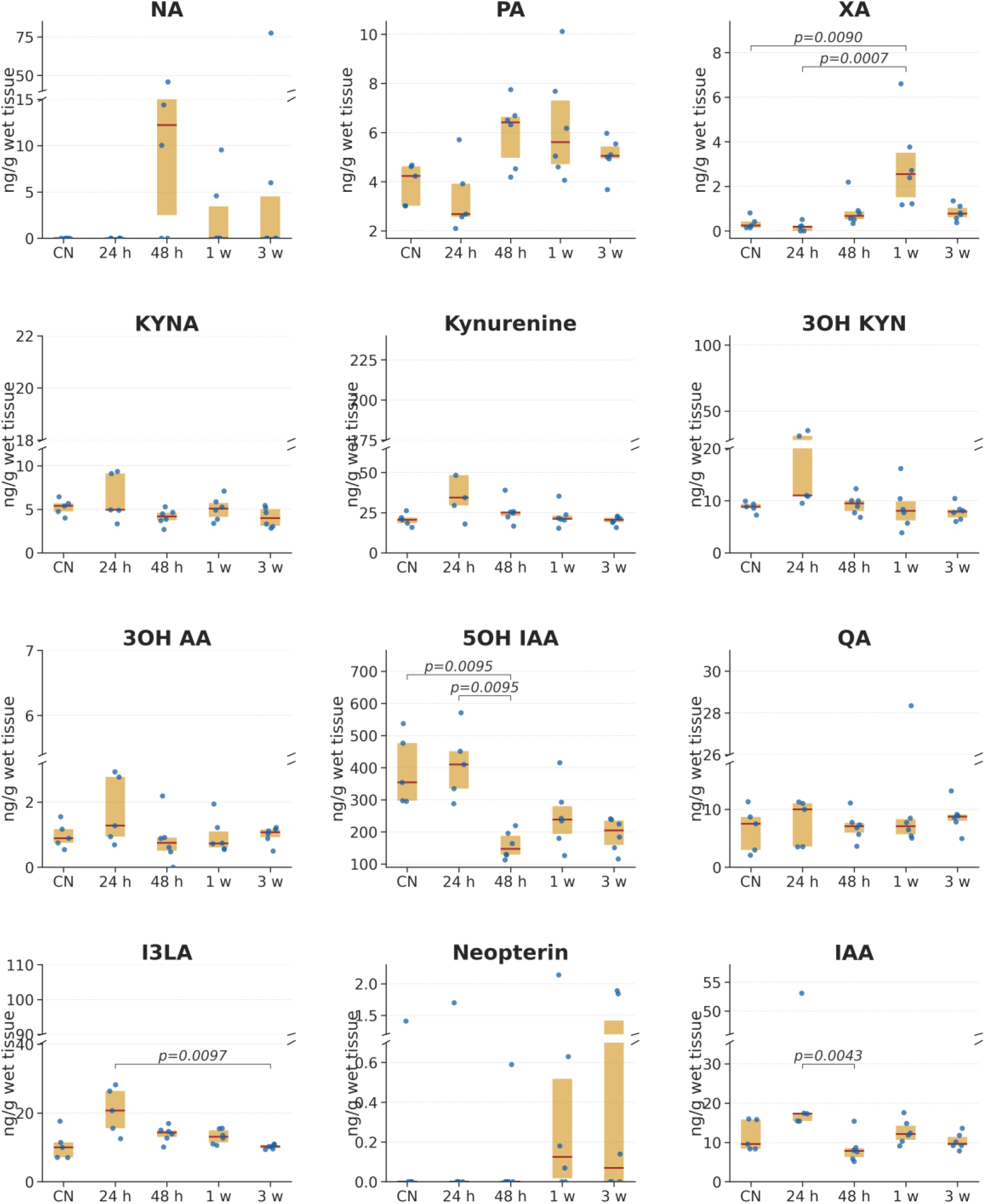
Changes of KP metabolites in frontal cortex after the HS. Four different time points were analyzed, 24h, 48h, 1 week and 3 weeks. Boxes represent the interquartile range with median (red line); individual data points are shown. p-values indicate significant pairwise differences (Dunn’s test with Bonferroni correction following Kruskal-Wallis). CN, control (n = 5); 24 h (n=5), 48 h, 1 w, 3 w: time points post-heatstroke (n = 6 per group).

5OH IAA was significantly reduced at 48 h post-HS compared with control animals (p = 0.001) and also showed a significant decrease relative to 24 h post-HS (p = 0.001), indicating an early suppression of serotonergic metabolism following thermal insult. A similar temporal reduction was observed for IAA, which was significantly decreased at 48 h compared with 24 h post-HS (p = 0.009). I3LA displayed a progressive declining trend over time, reaching a significant recovery at 3 weeks compared with 24 h post-HS (p = 0.008), suggesting sustained long-term modulation of indole metabolism in the frontal cortex.

In contrast, XA exhibited a delayed increase, with significantly elevated levels observed at 1 week compared with both control animals (p = 0.007) and the 24 h post-HS time point (p = 0.0007), indicating a time-dependent activation of this kynurenine branch during recovery. Both NA and PA showed a increased trend over time (ns). AA was below the limit of detection in brain tissue and was therefore not quantified.

## DISCUSSION

This is the first study to simultaneously quantify kynurenine-tryptophan pathway metabolites in both plasma and brain tissue (frontal cortex) in a mouse model of heatstroke. Using a fully validated LC-MS/MS method, we measured changes in 13 metabolites spanning the kynurenine, serotonin, and gut microbiota-derived indole branches of tryptophan metabolism.

Taken together, our findings reveal a biphasic metabolic response to HS, characterized by distinct early and late metabolic alterations (Fig. 5).

**Figure 5.**
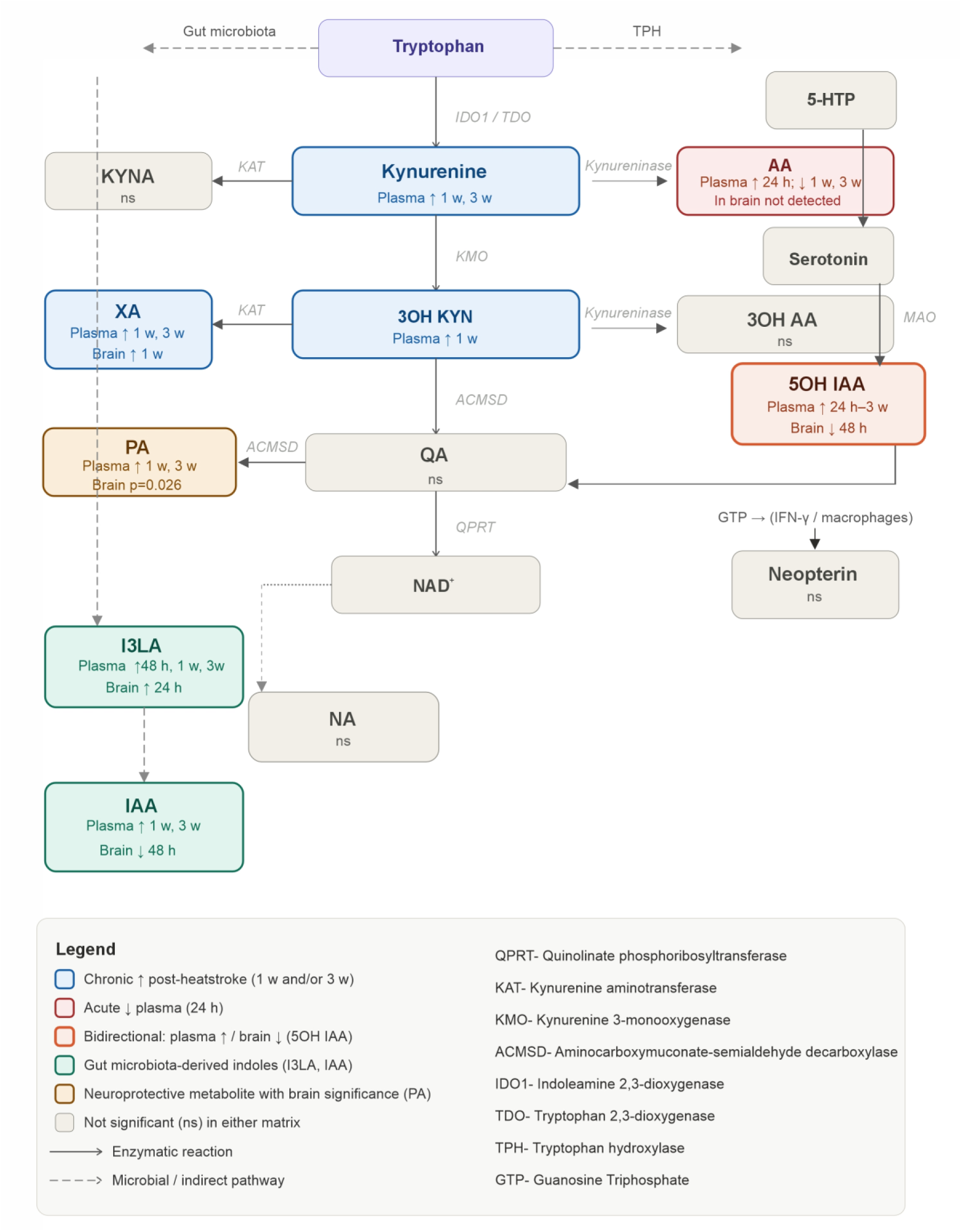
Schematic overwiev of KP changes detected in this study. Arrows indicate enzymatic reactions (solid) or microbial/indirect pathways (dashed).

We found progressive and persistent elevation of plasma kynurenine, 3-OH KYN, and XA at both 1 and 3 weeks following HS. These changes indicate prolonged activation of indoleamine 2,3-dioxygenase 1 (IDO1) and kynurenine 3-monooxygenase (KMO), two critical enzymes of the KP. IDO1, the rate-limiting enzyme of this pathway, is strongly induced by pro-inflammatory cytokines, particularly interferon-γ (IFN-γ) and tumour necrosis factor-α (TNF-α), which are known to remain elevated for several weeks after severe heat injury^30,31^. We suggest that the initial thermal and hypoxic insult initiates inflammatory response that outlasts the acute phase of HS by several weeks. This interpretation aligns with clinical observations in heatstroke survivors, in whom inflammatory biomarkers and neurocognitive impairments can persist for weeks to months after the acute event^32^. XA, a kynurenine aminotransferase (KAT)-derived metabolite of 3-OH KYN, was elevated in plasma at both 1 and 3 weeks and transiently increased in brain tissue at 1-week post-heatstroke. This pattern may reflect temporary disruption of the blood-brain barrier (BBB), facilitating the entry of peripheral kynurenine metabolites into the CNS before barrier integrity is restored by 3 weeks. XA has been reported to accumulate in several neuroinflammatory conditions and has been implicated in mitochondrial dysfunction and impaired insulin signalling at elevated concentrations^33^. Whether the increase in cerebral XA contributes to the neurological impairments associated with post-heatstroke recovery remains an important question for future investigation.

One of the most notable findings of our study is the opposing plasma-brain responses of 5OH IAA following HS. Plasma 5OH IAA concentrations remained elevated from 24 hours to 3 weeks post-insult, whereas brain 5OH IAA levels declined markedly, reaching their lowest levels at 48 hours. This pronounced plasma-brain dissociation suggests disruption of serotonergic homeostasis. Acute hyperthermia and autonomic dysregulation may initially trigger excessive serotonin release and turnover, reflected by the increase in circulating 5OH IAA. Subsequently, sustained KP activation may reduce tryptophan availability for serotonin biosynthesis, thereby contributing to depletion of central serotonergic stores. This phenomenon, commonly referred to as the “tryptophan steal” mechanism, has been described in sepsis and neuroinflammatory disorders ^34,35^. The present findings provide the first in vivo evidence supporting its involvement in heatstroke-induced metabolic dysfunction.

The implications of persistent serotonergic depletion may be substantial, given the central role of serotonin in regulating mood, thermoregulation, and sleep-wake homeostasis. Disruptions in these functions are frequently observed during post-heatstroke recovery^30^. Notably, brain 5OH IAA concentrations remained significantly reduced at both 1 and 3 weeks after heatstroke, indicating that serotonergic dysfunction persists well beyond the acute phase of injury. This sustained impairment suggests that restoration of serotonin metabolism may represent a promising therapeutic strategy during recovery.

Potential interventions could include supplementation with 5-hydroxytryptophan (5-HTP), a direct serotonin precursor, or treatment with selective serotonin reuptake inhibitors (SSRIs) to enhance serotonergic neurotransmission. Such approaches may be particularly relevant during the subacute recovery phase (48 h-1 week), when diversion of tryptophan metabolism toward the kynurenine pathway via IDO1 appears established, yet before chronic neuroinflammatory processes become fully entrenched. Nevertheless, further studies are needed to determine whether correction of serotonergic deficits can improve neurological, behavioural, and cognitive outcomes following heatstroke.

Plasma AA concentrations were significantly reduced 24 hours after HS and returned toward baseline by 48 hours. As AA is generated from kynurenine via kynureninase and has been reported to possess anti-inflammatory and antioxidant properties, its acute depletion may reflect preferential shunting of kynurenine metabolism toward the oxidative branch of the KP through kynurenine 3-monooxygenase (KMO), consistent with the concomitant increase in 3-OH KYN. This pattern suggests a transient shift toward the production of potentially neurotoxic and pro-oxidative metabolites during the acute phase of injury. Notably, AA was not detected in brain tissue in any experimental group. This observation is consistent with the low expression and activity of kynureninase within the CNS and the predominantly peripheral, particularly hepatic, origin of circulating AA^36^. The absence of measurable brain AA further supports the notion that peripheral and central kynurenine pathway metabolism are only partially coupled. Consequently, alterations in plasma metabolite concentrations may not directly reflect brain KP activity, which is determined primarily by local enzymatic regulation, and BBB permeability.

PA, a neuroprotective metabolite generated from QA through the action of aminocarboxymuconate semialdehyde decarboxylase, exhibited a strong trend toward significance in brain tissue, characterized by acute depletion at 24 hours followed by partial recovery at 48 hours. PA has been proposed to attenuate QA-mediated neurotoxicity through modulation of glutamatergic signaling and attenuation of NMDA receptor-dependent excitotoxicity, in contrast to QA, which acts as a potent NMDA receptor agonist^37^. The transient reduction in PA during the acute post-heatstroke phase may indicate diminished neuroprotective capacity at a time when inflammatory activation of the KP would be expected to favour QA production. This imbalance between neuroprotective and potentially neurotoxic metabolites could contribute to the enhanced vulnerability of the brain to excitotoxic and oxidative injury during the early stages of HS. The temporal association between reduced PA levels and the acute injury phase may therefore represent a mechanistic link to the period of maximal neurological dysfunction reported in both experimental models and clinical heatstroke.

IAA and I3LA are increasingly recognised as immunomodulatory metabolites that activate the aryl hydrocarbon receptor (AhR) and maintain intestinal immune homeostasis^38,39^. IAA and I3LA, two metabolites produced exclusively through bacterial metabolism of tryptophan, exhibited distinct temporal and compartment-specific alterations. Plasma IAA concentrations were significantly elevated at 1 and 3 weeks post-heatstroke, whereas brain IAA levels were significantly reduced at 48 hours. In contrast, I3LA showed no significant changes in plasma but was elevated in brain tissue at 24 hours. These findings suggest that HS may induce persistent alterations in gut microbial metabolism, consistent with evidence linking heat-related illness to intestinal barrier dysfunction, endotoxemia, and dysbiosis^40,41^. The sustained increase in circulating IAA may reflect enhanced microbial conversion of tryptophan to indole derivatives following intestinal injury and altered host-microbiome interactions. Conversely, the transient accumulation of I3LA within the brain may indicate heatstroke-induced disruption of BBB integrity during the acute phase. Together, these observations provide further evidence that HS affects not only host kynurenine metabolism but also the gut microbiota-brain axis, with distinct peripheral and central consequences that may contribute to both acute neurological injury and longer-term recovery processes. Restoration of gut microbiota diversity — for example through probiotic supplementation or dietary intervention — may represent a novel therapeutic target in HS recover.

To summarize our findings, the acute phase is characterized by AA depletion, marked peripheral serotonin activation, and reduced neuroprotective capacity. In contrast, the chronic phase is defined by sustained activation of the KP, depletion of central serotonin, and metabolic signatures consistent with gut microbiota dysbiosis. Among these alterations, the plasma-brain profile of 5OH IAA represents the most striking cross-compartment finding and may have important translational relevance.

## CONCLUSION

This study provides the first comprehensive characterization of kynurenine-tryptophan pathway metabolomics across both plasma and brain tissue in a mouse model of heatstroke, enabled by a fully validated simultaneous LC-MS/MS assay. The data reveal a biphasic metabolic response, characterized by an acute serotonergic dysregulation followed by chronic activation of the kynurenine pathway, with additional evidence suggesting that gut microbiota dysbiosis contributes to later-stage metabolic alterations. Among these findings, the plasma-brain dissociation of 5-hydroxyindoleacetic acid represents the most translationally relevant finding, offering a potential biomarker of central serotonergic depletion and supporting a rationale for serotonin-targeted interventions during the subacute recovery phase. Future studies incorporating region-specific brain analyses, direct measurements of brain serotonin and tryptophan and pharmacological intervention targeting IDO1 and KMO will be required to clarify the causal structure of these metabolic changes.

## AUTHOR CONTRIBUTIONS

DW, PM, and AK conceived and designed the study. AK and DW performed the in vivo experiments and collected the samples. PM, JP and AK conducted the LC-MS/MS measurements and data analyses. PM, DW, JP and AK revised the manuscript.

## PATIENT AND PUBLIC INVOLVEMENT

We have no patients or public involvement in the study herein.

## COMPETING INTERESTS

We have no competing interests in the study herein.

## ETHICS APPROVAL

All experiments on animals were carried out according to the institutional animal care guide-lines conforming to international standards and were approved by the State Veterinary and Food Committee of Slovak Republic.

## ACKNOWLEDGMENTS

This work was supported by EU NextGenerationEU through the Recovery and Resilience Plan for Slovakia under the project No. 09I03-03-V03-00086.

## List of Abbreviations

3OH AA: 3-hydroxyanthranilic acid
3OH KYN: 3-hydroxykynurenine
5-HTP: 5-hydroxytryptophan
5OH IAA: 5-hydroxyindole-3-acetic acid
AA: anthranilic acid
AADC: aromatic L-amino acid decarboxylase
ACMSD: aminocarboxymuconate semialdehyde decarboxylase
AhR: aryl hydrocarbon receptor
BBB: blood-brain barrier
CNS: central nervous system
CV: coefficient of variation
DMSO: dimethyl sulfoxide
EMA: European Medicines Agency
ESI: electrospray ionisation
FTN: flow-through needle
HS: heatstroke
HRI: heat-related illness
I3LA: indole-3-lactic acid
IAA: indole-3-acetic acid
IDO1: indoleamine 2,3-dioxygenase 1
IFN-γ: interferon-gamma
IS: internal standard
KAT: kynurenine aminotransferase
KMO: kynurenine 3-monooxygenase
KP: kynurenine pathway
KYNA: kynurenic acid
KYN: kynurenine
LC-MS/MS: liquid chromatography–tandem mass spectrometry
LOD: limit of detection
LOQ: limit of quantification
MAO: monoamine oxidase
MRM: multiple reaction monitoring
NA: nicotinic acid
NAD+: nicotinamide adenine dinucleotide
NMDA: N-methyl-D-aspartate
PA: picolinic acid
QA: quinolinic acid
QC: quality control
QPRT: quinolinate phosphoribosyltransferase
SRM: selected reaction monitoring
SSRI: selective serotonin reuptake inhibitor
TDO: tryptophan 2,3-dioxygenase
TNF-α: tumour necrosis factor-alpha
TPH: tryptophan hydroxylase
UPLC: ultra-performance liquid chromatography
XA: xanthurenic acid

## Notes

### Competing Interest Statement

The authors have declared no competing interest.

