## Supplementary Tables for "Kynurenine pathway metabolomics in heatstroke: a validated LC-MS/MS method reveals compartment-specific neurochemical disruption in a murine model": Majerova et al supplementary.docx

**Method validation**

| **Table S1.** Calibration parameters for LC-MS/MS quantification of 13 analytes in plasma. | | | | |
| --- | --- | --- | --- | --- |
| **Analyte** | **Regression equation** | **Linear range (ng/mL)** | **R²** | **LOQ (ng/mL)** |
| NA | *y = 0.78377x + 0.04968* | 0.1950–100.0 | 0.99966 | 0.1950 |
| PA | *y = 0.24185x - 0.00510* | 0.1950–100.0 | 0.99980 | 0.1950 |
| AA | *y = 0.34465x + 0.05138* | 0.1950–100.0 | 0.99920 | 0.1950 |
| KYNA | *y = 6.06913x + 0.69415* | 0.1950–100.0 | 0.99960 | 0.1950 |
| XA | *y = 0.21037x + 0.02171* | 0.3910–100.0 | 0.99763 | 0.3910 |
| Kynurenine | *y = 0.01296x - 0.00362* | 1.1720–600.0 | 0.99793 | 1.1720 |
| 3OH KYN | *y = 0.58600x + 0.04605* | 0.1950–100.0 | 0.99938 | 0.1950 |
| 3OH AA | *y = 0.06566x + 0.01360* | 0.1950–100.0 | 0.99955 | 0.1950 |
| 5OH IAA | *y = 2.45566x - 0.05916* | 0.1950–100.0 | 0.99935 | 0.1950 |
| QA | *y = 0.22907x - 0.01822* | 0.7810–400.0 | 0.99923 | 0.7810 |
| I3LA | *y = 0.24673x - 0.02785* | 0.7810–400.0 | 0.99922 | 0.7810 |
| IAA | *y = 0.23626x + 0.01535* | 0.7810–400.0 | 0.99995 | 0.7810 |
| Neopterin | *y = 0.11900x - 0.00352* | 0.1950–100.0 | 0.99934 | 0.1950 |

LOQ, lower limit of quantification (signal-to-noise ratio ≥10:1). R², coefficient of determination. Calibration curves constructed using 1/x weighted linear regression. y = response ratio (analyte peak area / internal standard peak area). KYNA, kynurenic acid; XA, xanthurenic acid; NA, nicotinic acid; PA, picolinic acid; AA, anthranilic acid; 3OH KYN, 3-hydroxykynurenine; 3OH AA, 3-hydroxyanthranilic acid; 5OH IAA, 5-hydroxyindole-3-acetic acid; QA, quinolinic acid; I3LA, indole-3-lactic acid; IAA, indole-3-acetic acid.

| **Table S2**. Calibration parameters for LC-MS/MS quantification of 13 analytes in brain tissue. | | | | |
| --- | --- | --- | --- | --- |
| **Analyte** | **Regression equation** | **Linear range (ng/mL)** | **R²** | **LOQ (ng/mL)** |
| NA | *y = 0.68814x + 2.58250* | 1.9500–500.0 | 0.99635 | 1.9500 |
| PA | *y = 0.31160x + 0.12281* | 0.0490–25.0 | 0.99578 | 0.0490 |
| AA | *y = 0.52566x + 0.20603* | 0.0980–25.0 | 0.99965 | 0.0980 |
| KYNA | *y = 5.44878x + 4.75529* | 0.0980–50.0 | 0.99990 | 0.0980 |
| XA | *y = 0.25900x + 0.06469* | 0.3910–12.5 | 0.99793 | 0.3910 |
| Kynurenine | *y = 0.00709x + 0.07459* | 3.1250–100.0 | 0.99994 | 3.1250 |
| 3OH KYN | *y = 0.57718x + 0.33463* | 0.1950–25.0 | 0.99975 | 0.1950 |
| 3OH AA | *y = 0.02731x + 0.01797* | 0.3910–12.5 | 0.99910 | 0.3910 |
| 5OH IAA | *y = 2.04116x + 130.74543* | 3.9000–500.0 | 0.99605 | 3.9000 |
| QA | *y = 0.24819x + 0.10151* | 0.1950–25.0 | 0.99720 | 0.1950 |
| I3LA | *y = 0.23398x + 0.20746* | 0.3900–25.0 | 0.99948 | 0.3900 |
| IAA | *y = 0.86648x + 0.30736* | 0.0490–25.0 | 0.99926 | 0.0490 |
| Neopterin | *y = 0.06286x + 0.00084* | 0.3910–12.5 | 0.99867 | 0.3910 |

LOQ, lower limit of quantification (signal-to-noise ratio ≥10:1). R², coefficient of determination. Calibration curves constructed using 1/x weighted linear regression. y = response ratio (analyte peak area / internal standard peak area). KYNA, kynurenic acid; XA, xanthurenic acid; NA, nicotinic acid; PA, picolinic acid; AA, anthranilic acid; 3OH KYN, 3-hydroxykynurenine; 3OH AA, 3-hydroxyanthranilic acid; 5OH IAA, 5-hydroxyindole-3-acetic acid; QA, quinolinic acid; I3LA, indole-3-lactic acid; IAA, indole-3-acetic acid.

| **Table S3.** Intraday accuracy and precision for LC-MS/MS quantification of 13 analytes in plasma. | | | | | | |
| --- | --- | --- | --- | --- | --- | --- |
| **Analyte** | **QC1** | | **QC2** | | **QC3** | |
|  | ***Acc (%)*** | ***CV (%)*** | ***Acc (%)*** | ***CV (%)*** | ***Acc (%)*** | ***CV (%)*** |
| NA | 102.1 | 4.4 | 95.9 | 4.3 | 97.2 | 2.9 |
| PA | 103.2 | 1.6 | 106.8 | 1.1 | 101.0 | 0.9 |
| AA | 101.4 | 4.3 | 96.5 | 1.0 | 96.7 | 1.9 |
| KYNA | 102.1 | 2.4 | 100.4 | 1.5 | 99.6 | 3.5 |
| XA | 102.6 | 4.3 | 98.8 | 1.6 | 100.9 | 2.7 |
| Kynurenine | 95.7 | 1.1 | 93.7 | 1.0 | 95.6 | 2.4 |
| 3OH KYN | 102.2 | 4.7 | 100.1 | 2.8 | 99.2 | 3.8 |
| 3OH AA | 95.8 | 3.5 | 97.4 | 1.0 | 99.0 | 0.6 |
| 5OH IAA | 104.4 | 2.3 | 100.0 | 1.8 | 99.0 | 2.2 |
| QA | 105.4 | 3.5 | 101.1 | 2.4 | 103.4 | 2.1 |
| I3LA | 103.4 | 1.3 | 102.8 | 1.9 | 104.9 | 1.1 |
| IAA | 105.3 | 1.3 | 104.1 | 2.1 | 104.9 | 1.6 |
| Neopterin | 103.2 | 4.6 | 102.1 | 2.1 | 100.4 | 1.8 |
| Acc, accuracy expressed as mean measured concentration / nominal concentration × 100%; CV, coefficient of variation. KYNA, kynurenic acid; XA, xanthurenic acid; NA, nicotinic acid; PA, picolinic acid; AA, anthranilic acid; 3OH KYN, 3-hydroxykynurenine; 3OH AA, 3-hydroxyanthranilic acid; 5OH IAA, 5-hydroxyindole-3-acetic acid; QA, quinolinic acid; I3LA, indole-3-lactic acid; IAA, indole-3-acetic acid. QC nominal concentrations (ng/mL) — QC1/QC2/QC3: NA, PA, AA, KYNA, XA, 3OH KYN, 3OH AA, 5OH IAA, Neopterin: 4/40/80; Kynurenine: 25/250/500; QA, I3LA, IAA: 15/150/300. n = 6 replicates per QC level. **Table S4.** Inter-day accuracy and precision for LC-MS/MS quantification of 13 analytes in plasma. \| \| \| \| \| \|; **Analyte** \| **QC1** \| \| **QC2** \| \| **QC3** \|; \| ***Acc (%)*** \| ***CV (%)*** \| ***Acc (%)*** \| ***CV (%)*** \| ***Acc (%)*** \| ***CV (%)***; NA \| 102.3 \| 3.6 \| 99.0 \| 4.2 \| 97.9 \| 3.4; PA \| 106.7 \| 5.5 \| 107.6 \| 4.1 \| 101.6 \| 2.9; AA \| 101.5 \| 3.2 \| 100.2 \| 2.8 \| 98.4 \| 1.9; KYNA \| 101.1 \| 3.4 \| 102.6 \| 2.3 \| 100.4 \| 2.6; XA \| 101.4 \| 3.8 \| 101.7 \| 3.4 \| 100.0 \| 3.3; Kynurenine \| 94.5 \| 2.5 \| 96.2 \| 3.3 \| 98.5 \| 3.1; 3OH KYN \| 102.6 \| 3.1 \| 102.0 \| 3.5 \| 98.9 \| 4.6; 3OH AA \| 99.6 \| 4.1 \| 101.6 \| 3.3 \| 98.8 \| 1.7; 5OH IAA \| 103.8 \| 2.4 \| 101.7 \| 2.0 \| 100.2 \| 2.2; QA \| 105.8 \| 3.4 \| 105.4 \| 4.9 \| 106.0 \| 3.5; I3LA \| 105.5 \| 2.5 \| 105.6 \| 3.7 \| 106.1 \| 2.2; IAA \| 107.2 \| 5.0 \| 107.2 \| 4.2 \| 104.8 \| 2.3; Neopterin \| 101.1 \| 4.3 \| 98.2 \| 5.5 \| 98.5 \| 3.5; Acc, accuracy expressed as mean measured concentration / nominal concentration × 100%; CV, coefficient of variation. KYNA, kynurenic acid; XA, xanthurenic acid; NA, nicotinic acid; PA, picolinic acid; AA, anthranilic acid; 3OH KYN, 3-hydroxykynurenine; 3OH AA, 3-hydroxyanthranilic acid; 5OH IAA, 5-hydroxyindole-3-acetic acid; QA, quinolinic acid; I3LA, indole-3-lactic acid; IAA, indole-3-acetic acid.QC nominal concentrations (ng/mL) — QC1/QC2/QC3: NA, PA, AA, KYNA, XA, 3OH KYN, 3OH AA, 5OH IAA, Neopterin: 4/40/80; Kynurenine: 25/250/500; QA, I3LA, IAA: 15/150/300. All compounds validated across three independent analytical runs (n = 18 per QC level) (**Table S5**. Intraday accuracy and precision for LC-MS/MS quantification of 13 analytes in brain tissue . \| \| \| \| \| \|; **Analyte** \| **QC1** \| \| **QC2** \| \| **QC3** \|; \| ***Acc (%)*** \| ***CV (%)*** \| ***Acc (%)*** \| ***CV (%)*** \| ***Acc (%)*** \| ***CV (%)***; NA \| 104.8 \| 2.9 \| 105.5 \| 4.9 \| 102.9 \| 3.0; PA \| 96.3 \| 12.4 \| 99.0 \| 8.6 \| 102.2 \| 2.6; AA \| 89.3 \| 4.9 \| 96.1 \| 1.6 \| 99.5 \| 2.0; KYNA \| 91.1 \| 6.3 \| 92.2 \| 2.4 \| 96.6 \| 2.1; XA \| 97.6 \| 7.5 \| 90.7 \| 3.8 \| 100.4 \| 1.7; Kynurenine \| 113.2 \| 4.8 \| 102.3 \| 3.4 \| 103.0 \| 2.0; 3OH KYN \| 88.8 \| 9.5 \| 95.0 \| 3.2 \| 99.8 \| 2.5; 3OH AA \| 95.2 \| 5.0 \| 102.8 \| 6.7 \| 108.7 \| 2.9; 5OH IAA \| 106.4 \| 4.4 \| 98.7 \| 2.0 \| 99.6 \| 2.9; QA \| 108.4 \| 6.2 \| 97.2 \| 5.0 \| 100.6 \| 5.3; I3LA \| 101.6 \| 4.9 \| 95.5 \| 3.1 \| 101.6 \| 1.5; IAA \| 99.9 \| 2.1 \| 98.7 \| 1.7 \| 100.0 \| 2.4; Neopterin \| 104.5 \| 8.5 \| 98.7 \| 5.7 \| 101.1 \| 3.6; Acc, accuracy = mean measured / nominal × 100%; CV, coefficient of variation. Acceptance criteria: Acc 85–115%, CV ≤15%. KYNA, kynurenic acid; XA, xanthurenic acid; NA, nicotinic acid; PA, picolinic acid; AA, anthranilic acid; 3OH KYN, 3-hydroxykynurenine; 3OH AA, 3-hydroxyanthranilic acid; 5OH IAA, 5-hydroxyindole-3-acetic acid; QA, quinolinic acid; I3LA, indole-3-lactic acid; IAA, indole-3-acetic acid. QC nominal concentrations (ng/mL) — QC1/QC2/QC3: NA, 5OH IAA: 25/100/400; Kynurenine: 5/20/80; PA, AA, 3OH KYN, QA, I3LA, IAA: 1.25/5/20; KYNA: 2.5/10/40; XA, 3OH AA, Neopterin: 0.625/2.5/10. n = 6 replicates per QC level. (**Table S6**. Inter-day accuracy and precision for LC-MS/MS quantification of 13 analytes in brain tissue. \| \| \| \| \| \| \|; **Analyte** \| **n** \| **QC1** \| \| **QC2** \| \| **QC3** \|; \| \| ***Acc (%)*** \| ***CV (%)*** \| ***Acc (%)*** \| ***CV (%)*** \| ***Acc (%)*** \| ***CV (%)***; NA \| 18 \| 102.7 \| 2.7 \| 104.5 \| 3.3 \| 101.7 \| 2.2; PA \| 18 \| 101.9 \| 7.6 \| 101.8 \| 5.2 \| 100.1 \| 2.7; AA \| 18 \| 95.7 \| 6.0 \| 98.6 \| 2.3 \| 99.0 \| 1.3; KYNA \| 18 \| 95.4 \| 5.3 \| 97.0 \| 3.9 \| 97.1 \| 1.4; XA \| 18 \| 98.1 \| 5.6 \| 94.3 \| 4.4 \| 100.0 \| 1.7; Kynurenine \| 18 \| 102.9 \| 7.9 \| 101.8 \| 2.4 \| 102.7 \| 1.2; 3OH KYN \| 18 \| 101.6 \| 10.7 \| 100.0 \| 4.3 \| 99.7 \| 1.8; 3OH AA \| 18 \| 96.2 \| 4.3 \| 103.8 \| 4.1 \| 102.5 \| 8.3; 5OH IAA \| 18 \| 102.0 \| 5.4 \| 100.0 \| 2.3 \| 100.5 \| 2.2; QA \| 18 \| 101.7 \| 6.2 \| 96.7 \| 4.1 \| 97.6 \| 4.1; I3LA \| 18 \| 99.1 \| 3.9 \| 99.5 \| 3.7 \| 99.5 \| 2.0; IAA \| 18 \| 101.4 \| 2.6 \| 101.8 \| 2.5 \| 100.0 \| 1.7; Neopterin \| 18 \| 98.3 \| 8.3 \| 99.6 \| 5.1 \| 100.8 \| 2.3; Acc, accuracy = mean measured / nominal × 100%; CV, coefficient of variation. Acceptance criteria: Acc 85–115%, CV ≤15%. KYNA, kynurenic acid; XA, xanthurenic acid; NA, nicotinic acid; PA, picolinic acid; AA, anthranilic acid; 3OH KYN, 3-hydroxykynurenine; 3OH AA, 3-hydroxyanthranilic acid; 5OH IAA, 5-hydroxyindole-3-acetic acid; QA, quinolinic acid; I3LA, indole-3-lactic acid; IAA, indole-3-acetic acid. QC nominal concentrations (ng/mL) — QC1/QC2/QC3: NA, 5OH IAA: 25/100/400; Kynurenine: 5/20/80; PA, AA, 3OH KYN, QA, I3LA, IAA: 1.25/5/20; KYNA: 2.5/10/40; XA, 3OH AA, Neopterin: 0.625/2.5/10. Data from three independent analytical runs (n = 18). \| \| \| \| \| \| \|) \| \| \| \| \| \|) \| \| \| \| \| \| | | | | | | |

| **Table S7**. Extraction recovery and matrix effect for LC-MS/MS quantification of 13 analytes in brain tissue. | | | | |
| --- | --- | --- | --- | --- |
| **Analyte** | **QC1** | | **QC2** | |
|  | ***Rec (%)*** | ***ME (%)*** | ***Rec (%)*** | ***ME (%)*** |
| NA | 83.8 | 100.1 | 84.3 | 99.5 |
| PA | 98.9 | 92.0 | 99.2 | 98.7 |
| AA | 91.1 | 107.8 | 90.8 | 120.0 |
| KYNA | 99.0 | 101.2 | 90.2 | 105.0 |
| XA | 94.6 | 103.1 | 87.9 | 112.7 |
| Kynurenine | 99.2 | 100.8 | 104.0 | 97.0 |
| 3OH KYN | 94.9 | 100.0 | 87.4 | 99.0 |
| 3OH AA | 93.9 | 113.9 | 88.6 | 115.7 |
| 5OH IAA | 100.7 | 98.5 | 97.8 | 104.0 |
| QA | 95.5 | 115.9 | 92.8 | 113.5 |
| I3LA | 99.9 | 103.7 | 95.9 | 111.6 |
| IAA | 97.0 | 100.7 | 99.0 | 78.5 |
| Neopterin | 83.2 | 105.6 | 85.0 | 113.2 |
| Rec, extraction recovery = analyte response after extraction relative to a post-extraction spike at the same nominal level. ME, matrix effect = analyte response in post-extraction spiked matrix relative to neat solution. KYNA, kynurenic acid; XA, xanthurenic acid; NA, nicotinic acid; PA, picolinic acid; AA, anthranilic acid; 3OH KYN, 3-hydroxykynurenine; 3OH AA, 3-hydroxyanthranilic acid; 5OH IAA, 5-hydroxyindole-3-acetic acid; QA, quinolinic acid; I3LA, indole-3-lactic acid; IAA, indole-3-acetic acid. QC1/2 nominal spike concentrations (ng/mL): Kynurenine, QA, I3LA, 5/40; all other analytes, 1.25/10. | | | | |

| **Table S8**. Autosampler stability of 13 analytes (stored at 10°C for 24 h). | | | | | | | | | | |
| --- | --- | --- | --- | --- | --- | --- | --- | --- | --- | --- |
| **Analyte** | **QC1** | | | **QC2** | | | | **QC3** | | |
|  | ***Mean (ng/mL)*** | ***Stab (%)*** | | ***Mean (ng/mL)*** | | ***Stab (%)*** | | ***Mean (ng/mL)*** | | ***Stab (%)*** |
| NA | 4.10 | 102.4 | | 40.58 | | 101.4 | | 77.28 | | 96.6 |
| PA | 4.13 | 103.2 | | 42.73 | | 106.8 | | 81.08 | | 101.3 |
| AA | 4.15 | 103.7 | | 41.06 | | 102.7 | | 79.17 | | 99.0 |
| KYNA | 4.12 | 103.0 | | 41.64 | | 104.1 | | 79.49 | | 99.4 |
| XA | 4.00 | 100.0 | | 40.62 | | 101.6 | | 77.51 | | 96.9 |
| Kynurenine | 22.96 | 91.9 | | 236.17 | | 94.5 | | 490.66 | | 98.1 |
| 3OH KYN | 4.09 | 102.2 | | 40.21 | | 100.5 | | 75.69 | | 94.6 |
| 3OH AA | 4.13 | 103.3 | | 41.70 | | 104.2 | | 78.06 | | 97.6 |
| 5OH IAA | 4.20 | 105.0 | | 41.22 | | 103.0 | | 79.43 | | 99.3 |
| QA | 16.24 | 108.3 | | 167.59 | | 111.7 | | 329.79 | | 109.9 |
| I3LA | 16.25 | 108.3 | | 161.03 | | 107.4 | | 315.29 | | 105.1 |
| IAA | 17.09 | 113.9 | | 169.47 | | 113.0 | | 318.49 | | 106.2 |
| Neopterin | 4.13 | 103.2 | | 40.86 | | 102.1 | | 80.31 | | 100.4 |
| KYNA, kynurenic acid; XA, xanthurenic acid; NA, nicotinic acid; PA, picolinic acid; AA, anthranilic acid; 3OH KYN, 3-hydroxykynurenine; 3OH AA, 3-hydroxyanthranilic acid; 5OH IAA, 5-hydroxyindole-3-acetic acid; QA, quinolinic acid; I3LA, indole-3-lactic acid; IAA, indole-3-acetic acid.QC nominal concentrations (ng/mL) — QC1/QC2/QC3: NA, PA, AA, KYNA, XA, 3OH KYN, 3OH AA, 5OH IAA, Neopterin: 4/40/80; Kynurenine: 25/250/500; QA, I3LA, IAA: 15/150/300. n = 6 replicates per QC level. Samples stored at 10°C in the autosampler for 24 h prior to re-analysis. | | | | | | | | | | |

| **Table S9.** Freeze–thaw stability of 13 analytes in plasma after three freeze–thaw cycles (−80°C/room temperature). | | | | | | |
| --- | --- | --- | --- | --- | --- | --- |
| **Analyte** | **QC1** | | **QC2** | | **QC3** | |
|  | ***Mean (ng/mL)*** | ***% of nominal*** | ***Mean (ng/mL)*** | ***% of nominal*** | ***Mean (ng/mL)*** | ***% of nominal*** |
| NA | 4.13 | 103.3 | 40.49 | 101.2 | 76.27 | 95.3 |
| PA | 4.27 | 106.8 | 41.62 | 104.1 | 81.87 | 102.3 |
| AA | 4.09 | 102.2 | 41.02 | 102.5 | 78.52 | 98.2 |
| KYNA | 4.08 | 102.1 | 41.26 | 103.1 | 77.95 | 97.4 |
| XA | 3.98 | 99.4 | 40.78 | 102.0 | 80.22 | 100.3 |
| Kynurenine | 22.92 | 91.7 | 235.57 | 94.2 | 485.93 | 97.2 |
| 3OH KYN | 4.03 | 100.8 | 41.10 | 102.7 | 77.49 | 96.9 |
| 3OH AA | 4.22 | 105.5 | 41.36 | 103.4 | 78.92 | 98.7 |
| 5OH IAA | 4.16 | 104.0 | 41.44 | 103.6 | 80.62 | 100.8 |
| QA | 16.41 | 109.4 | 169.79 | 113.2 | 330.20 | 110.1 |
| I3LA | 16.31 | 108.7 | 160.90 | 107.3 | 314.97 | 105.0 |
| IAA | 17.26 | 115.0 | 168.91 | 112.6 | 319.67 | 106.6 |
| Neopterin | 4.19 | 104.9 | 41.40 | 103.5 | 79.46 | 99.3 |
| % of nominal, mean measured concentration / nominal concentration × 100%. KYNA, kynurenic acid; XA, xanthurenic acid; NA, nicotinic acid; PA, picolinic acid; AA, anthranilic acid; 3OH KYN, 3-hydroxykynurenine; 3OH AA, 3-hydroxyanthranilic acid; 5OH IAA, 5-hydroxyindole-3-acetic acid; QA, quinolinic acid; I3LA, indole-3-lactic acid; IAA, indole-3-acetic acid. QC nominal concentrations (ng/mL) — QC1/QC2/QC: NA, PA, AA, KYNA, XA, 3OH KYN, 3OH AA, 5OH IAA, Neopterin: 4/40/80; Kynurenine: 25/250/500; QA, I3LA, IAA: 15/150/300. Samples subjected to three freeze–thaw cycles (frozen at −80°C, thawed at room temperature) prior to analysis. n = 6 replicates per QC level. **Table S10**. Freeze–thaw stability of 13 analytes in brain tissue after three freeze–thaw cycles (−80°C/room temperature). \| \| \| \| \| \| \| \| \| \| \| \| \|; **Analyte** \| **QC1** \| \| \| \| **QC2** \| \| \| \| **QC3** \| \| \| \|; \| ***Mean (ng/mL)*** \| \| ***Stab (%)*** \| \| ***Mean (ng/mL)*** \| \| ***Stab (%)*** \| \| ***Mean (ng/mL)*** \| \| ***Stab (%)*** \| \|; NA \| 25.75 \| \| 103.0 \| \| 105.52 \| \| 105.5 \| \| 407.46 \| \| 101.9 \| \|; PA \| 1.33 \| \| 106.4 \| \| 5.01 \| \| 100.2 \| \| 19.86 \| \| 99.3 \| \|; AA \| 1.19 \| \| 95.2 \| \| 4.87 \| \| 97.4 \| \| 19.35 \| \| 96.7 \| \|; KYNA \| 2.15 \| \| 85.8 \| \| 8.63 \| \| 86.3 \| \| 34.71 \| \| 86.8 \| \|; XA \| 0.60 \| \| 95.7 \| \| 2.29 \| \| 91.5 \| \| 8.93 \| \| 89.3 \| \|; Kynurenine \| 5.18 \| \| 103.6 \| \| 21.46 \| \| 107.3 \| \| 82.19 \| \| 102.7 \| \|; 3OH KYN \| 1.19 \| \| 95.1 \| \| 4.88 \| \| 97.6 \| \| 19.44 \| \| 97.2 \| \|; 3OH AA \| 0.87 \| \| 139.5 \| \| 3.19 \| \| 127.7 \| \| 13.99 \| \| 139.9 \| \|; 5OH IAA \| 21.44 \| \| 85.7 \| \| 98.48 \| \| 98.5 \| \| 381.65 \| \| 95.4 \| \|; QA \| 1.25 \| \| 100.0 \| \| 5.00 \| \| 100.0 \| \| 18.65 \| \| 93.3 \| \|; I3LA \| 1.17 \| \| 93.9 \| \| 4.84 \| \| 96.9 \| \| 19.29 \| \| 96.4 \| \|; IAA \| 1.23 \| \| 98.3 \| \| 5.06 \| \| 101.2 \| \| 19.69 \| \| 98.5 \| \|; Neopterin \| 0.61 \| \| 98.4 \| \| 2.54 \| \| 101.5 \| \| 10.07 \| \| 100.7 \| \|; Stab, stability = mean measured / nominal × 100%. Acceptance criteria: Stab 85–115%. Values in red bold indicate failure. KYNA, kynurenic acid; XA, xanthurenic acid; NA, nicotinic acid; PA, picolinic acid; AA, anthranilic acid; 3OH KYN, 3-hydroxykynurenine; 3OH AA, 3-hydroxyanthranilic acid; 5OH IAA, 5-hydroxyindole-3-acetic acid; QA, quinolinic acid; I3LA, indole-3-lactic acid; IAA, indole-3-acetic acid.QC nominal concentrations (ng/mL) — QC1/QC2/QC3: NA, 5OH IAA: 25/100/400; Kynurenine: 5/20/80; PA, AA, 3OH KYN, QA, I3LA, IAA: 1.25/5/20; KYNA: 2.5/10/40; XA, 3OH AA, Neopterin: 0.625/2.5/10. n = 6 per QC level. \| \| \| \| \| \| \| \| \| \| \| \| \| | | | | | | |
